# Deep Learning Identification of Stiffness Markers in Breast Cancer

**DOI:** 10.1101/2020.12.17.423077

**Authors:** Alexandra Sneider, Joo Ho Kim, Ashley Kiemen, Pei-Hsun Wu, Mehran Habibi, Marissa White, Jude M. Phillip, Luo Gu, Denis Wirtz

## Abstract

While essential to our understanding of solid tumor progression, the study of cell and tissue mechanics has yet to find traction in the clinic. Determining tissue stiffness, a mechanical property known to promote a malignant phenotype *in vitro* and *in vivo*, is not part of the standard algorithm for the diagnosis and treatment of breast cancer. Instead, clinicians routinely use mammograms to identify malignant lesions and radiographically dense breast tissue is associated with an increased risk of developing cancer. Whether breast density is related to tumor tissue stiffness, and what cellular and non-cellular components of the tumor contribute the most to its stiffness are not well understood. Through training of a deep learning network and mechanical measurements of fresh patient tissue, we create a bridge in understanding between clinical and mechanical markers. The automatic identification of cellular and extracellular features from hematoxylin and eosin (H&E)-stained slides reveals that global and local breast tissue stiffness best correlate with the percentage of straight collagen. Global breast tissue mechanics correlate weakly with the percentage of blood vessels and fibrotic tissue, and non-significantly with the percentage of fat, ducts, tumor cells, and wavy collagen in tissue. Importantly, the percentage of dense breast tissue does not directly correlate with tissue stiffness or straight collagen content.

## Introduction

A significant disconnect exists between sophisticated biomechanical and biophysical experiments “at the bench,”*(1)* and clinical methods used to determine effective therapeutics for patients with solid tumors. Women with breast cancer are typically diagnosed via dedicated breast imaging modalities (mammogram, ultrasound, MRI, tomosynthesis). Mammograms are radiological images that reveal regions of dense, fibrous, and glandular breast tissue typically shown in white against non-dense, fatty tissue in black.*(2)* Methods for evaluating breast density include visually binning images into categories (fatty, scattered, heterogenous, extremely dense) based on the percentage of white versus black features in the breast image, or quantifying the exact percentage of dense tissue in white via image analysis (Fig. 1a).

**Fig. 1.**
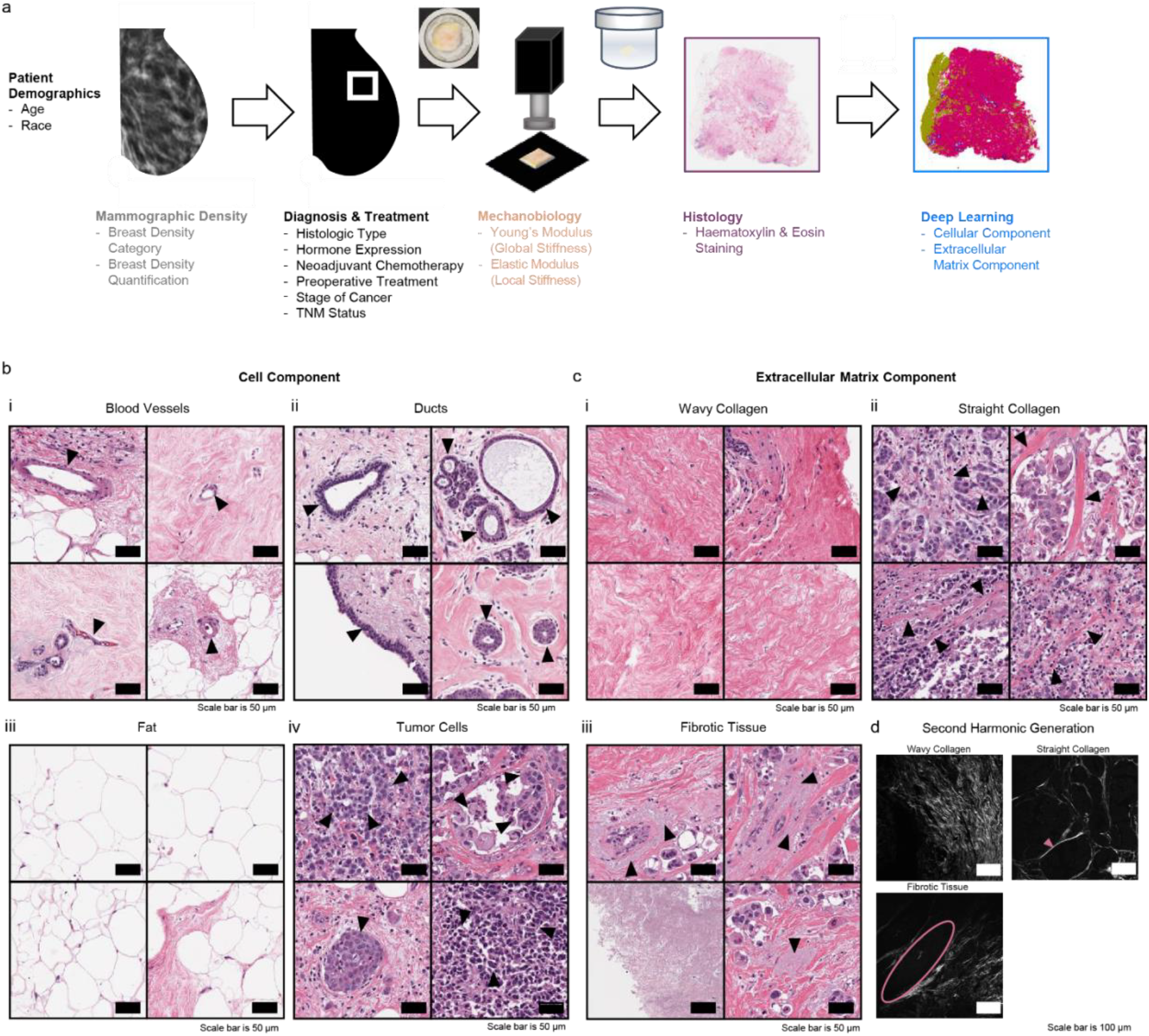
Breast tissue acquisition, characterization, and selected classes for deep learning composition analysis. **A**, Schematic detailing the breast tissue acquisition and characterization starting with medical imaging via mammogram, diagnosis, treatment, mechanical measurements, histology, and machine learning. **B**, Hematoxylin and eosin (H&E)-stained images of cell component classes including (**i**) blood vessels (capillaries, venules/arterioles), (**ii**) ducts (excretory, terminal/acini/alveoli), (**iii**) fat, (**iv**) tumor cells. Scale bars in black are 50 μm. **C**, Hematoxylin and eosin (H&E)-stained images of extracellular matrix component (ECM) classes including (**i**) wavy collagen, (**ii**) straight collagen, and (**iii**) fibrotic tissue. Scale bars in black are 50 μm. **D**, Second harmonic generation (SHG) images confirming (**i**) the wavy ECM class is wavy collagen, (**ii**) the straight ECM class is straight collagen, and (**iii**) the fibrotic tissue is not collagen detectable with SHG. Scale bars in white are 100 μm.

Dense breast tissue poses two major risks for patients. The first is an impaired ability to detect malignant lesions through imaging.*(3)* The second is as an independent risk factor for breast cancer. Increased breast density is associated with a worse patient prognosis,*(4–13)* poor progression free survival rate,*(14, 15)* and increased mortality.*(16, 17)* These denser tissue regions are purported to be more fibrous than the surrounding tissue,*(18)* and have been linked to an increase in the amount of collagen and numbers of epithelial and non-epithelial cells.*(19)*

While mammography remains the standard for breast cancer screening, other imaging methods like elastography have been developed to leverage changes in tissue stiffness.*(20–23)* Breast ultrasound elastography, a method utilizing sonographic imaging, identifies changes in elastic moduli to detect lesions in the breast*(24, 25)* and shows promise as an imaging modality alongside traditional ultrasound or mammograms to further characterize masses.*(26, 27)* After using multiple imaging modalities, core needle biopsies are still an essential next step in the diagnostic algorithm.*(28)*

In the laboratory, the application of cell and tissue mechanics has yielded great insight into tumor development and progression.*(29–40)* Tissue stiffening, widely attributed to an increase in collagen deposition and cross-linking,*(41–44)* has been proposed as a marker of tumor biogenesis. Recent studies assessing mechanical tissue stiffness often use previously frozen or fixed samples;*(44–46)* however these preservation processes significantly impact the resulting mechanical measurements.*(47)* Despite the lack of a direct link, many conflate breast tissue density (radiographically defined fibrous and glandular tissue) and breast tissue stiffness (the resistance of tissue to deformation;*(48)* often broadly referring to the elastic modulus). The disconnect in terminology, between breast density *vs*. breast stiffness, and assessed features in the clinic *vs.* the bench significantly hampers the generation of new and effective mechanobiology-inspired cancer therapies.*(49–53)*

Here, we relate patient information, medical imaging, treatment history, and histology to global and local mechanical measurements through the use of a deep learning, convolutional neural network (CNN) that accurately identifies tissue components from hematoxylin and eosin (H&E)-stained sections of breast cancer tissues (Fig. 1a). Patients with luminal A subtype, estrogen receptor (ER) and/or progesterone receptor (PR) positive and HER2 negative, have dense breasts that have been linked to an increased breast cancer risk.*(54)* Patients with triple-negative (TNBC) subtype (ER, PR, HER2 negative) tend to have lower mammographic breast density than non-TNBC patients.*(55, 56)* Herein we utilize 32 tissue samples from nine patients with a luminal A subtype and one patient with a triple-negative (ER, PR, HER2 negative) subtype. Global stiffness is determined by a compression test, which consists of taking one uniaxial measurement per tissue sample to obtain Young’s modulus. Local stiffness, obtained through microindentation, reports the elastic modulus from multiple, evenly spaced indentation measurements across the same tissue surface. Based on these measurements, we then identify correlations between tissue stiffness, tissue composition, and breast density.

## Results

### Deep-learning model classifies essential cellular and extracellular matrix features

Patients received diagnostic breast imaging via mammogram, pathologic examination, and characterization, and finally surgery prior to release of tissue samples for mechanical measurements, H&E staining, and deep learning analysis (Fig. 1a, See Methods). This study presents analysis of ten patients, with stiffness measurements on samples from nine of the ten patients (Table 1).

**Table 1.**
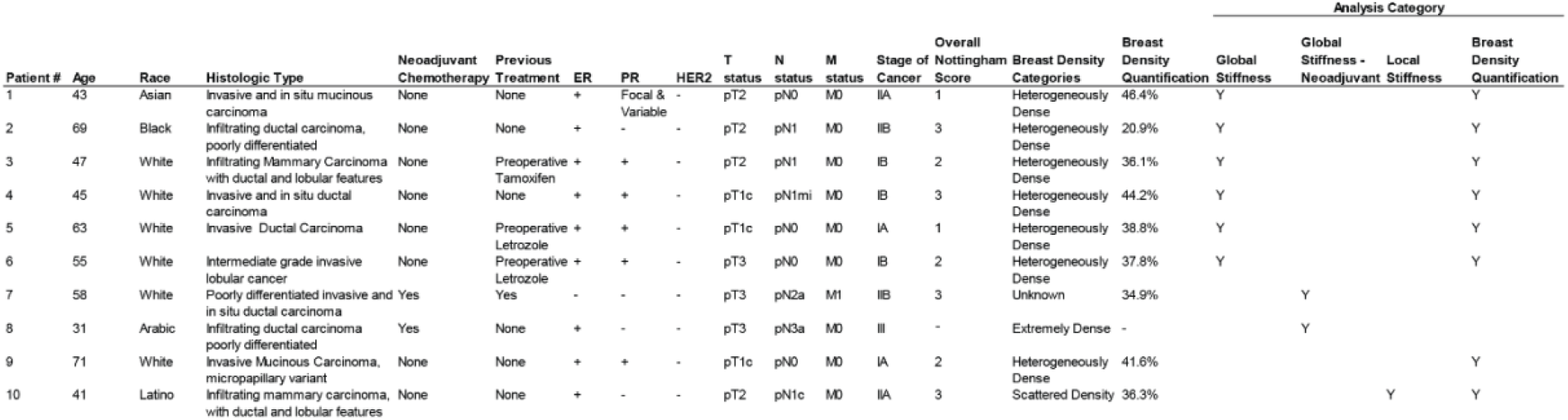
Patient cohort demographic and pathologic information.

Breast tissue histology is complex and heterogeneous, as many components change in content and organization during tumor progression.*(57)* Deep learning classifiers have been successful in identifying normal and cancerous components in histological sections.*(58, 59)* This paper utilizes a CNN-based deep learning pipeline which has previously shown success in classification of histological images into pathologically relevant subtypes.*(60)* We identified seven clinically relevant and computationally identifiable tissue classes consistent across most tested breast tissues (Fig. 1b and 1c). The four cell component classes are blood vessels (capillaries and venules/arterioles), ducts (excretory, terminal/acini/alveoli), fat, and tumor cells (viable, necrotic) (Fig. 1b). The three extracellular matrix (ECM) classes are wavy collagen, straight collagen, and fibrotic tissue (Fig. 1c). Second harmonic generation confirmed that the wavy and straight ECM classes were fibrillar collagen (Fig. 1d). The wavy and straight stromal phenotypes, a distinction which has been noted by others,*(61)* were identified from a visual assessment of the histology sections. Our eighth class, white space, encapsulates all non-tissue space on the images (not shown).

The CNN successfully identified and classified the seven cell and tissue classes stated above in 32 patient tissue samples consisting of 13 tumor-adjacent and 19 tumor samples from all ten patients (Fig. 2a). The confusion matrix details class accuracy in the testing dataset (Fig. 2b). Overall testing accuracy was 93.0% (Fig. 2b). All tissue classes were identified with greater than 90% sensitivity, except for fat cells at 89.7%. In this case, fat tended to be misclassified as white space due to the chosen image window size in the neural net. Histological subtyping revealed that a subset of luminal A tumors has ductal morphologies, which could explain why ducts and tumor cells were misclassified as each other 2.5% of the time (Fig. 2b). Wavy collagen was misclassified as straight collagen 3.2% of the time, however, straight collagen was never mistaken for wavy collagen (Fig. 2b). The successful separation of these ECM phenotypes was important for ensuring that we could analyze the contribution of the stroma to global and local modulus measurements. Any incorrectly classified straight collagen tended to be attributed to the tumor cell class, which was most likely a biological result of short straight fibers amongst tumor cells. White space misclassified as other cellular classes may be due to the presence of lumen (Fig. 2b). Visual comparison highlights the trained network’s ability to distinguish histological features even in complex tissue microenvironments (Fig. 2c and Fig. S1a).

**Fig. 2.**
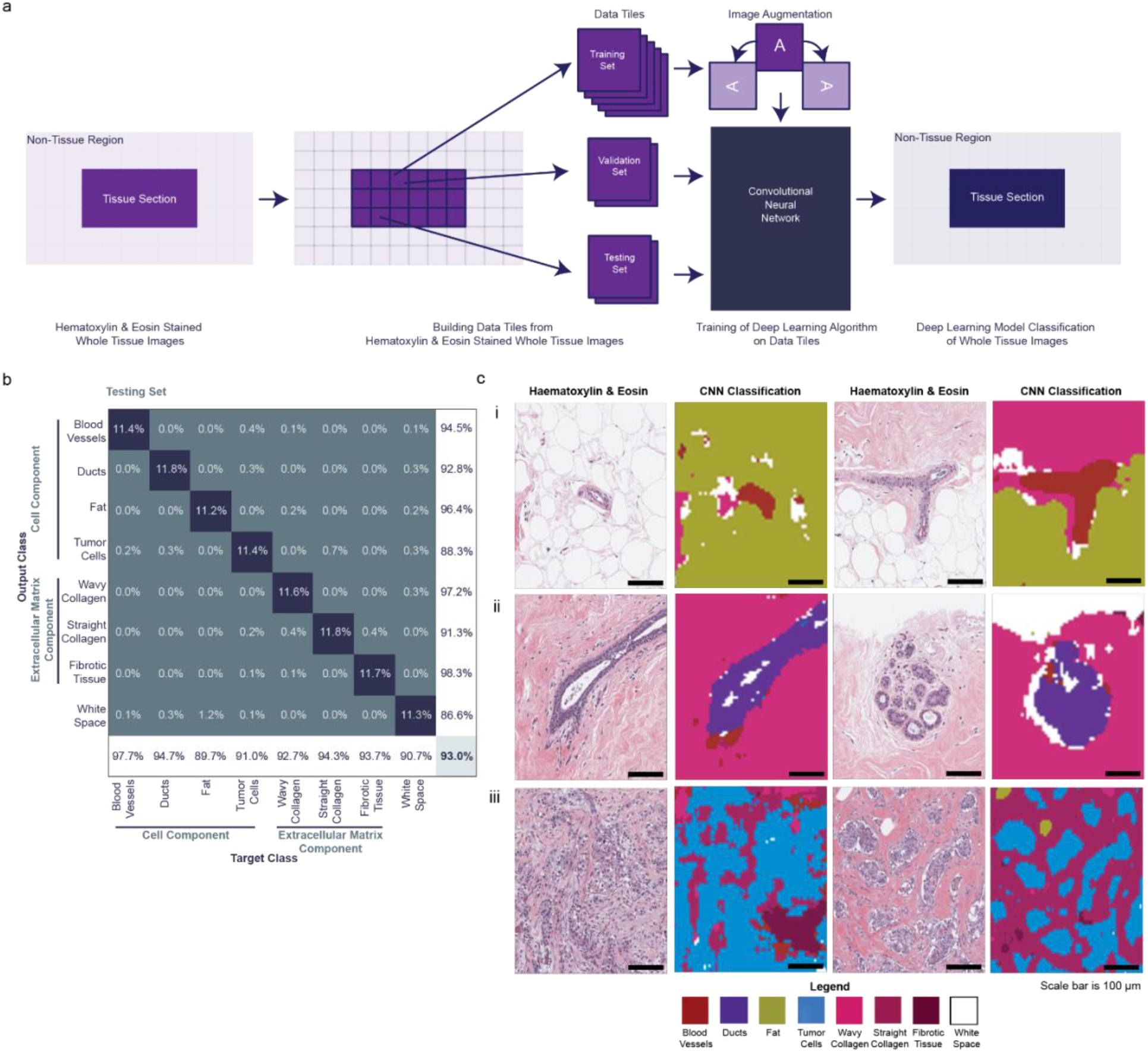
Convolutional Neural Network construction, quantitative and qualitative analysis. (**A**) Schematic showing the division of H&E stained tissue slides (32 tissues, 10 patients) into data tiles for training, validation, and testing. While each dataset is from the same patient tissue slides, the testing set was developed from a separate set of annotations than the training and validation sets. The training images are augmented by rotation [−90°,90°] before use in the convolutional neural network (CNN). The accuracy of the CNN is determined against the testing sets. Finally, the whole tissue images are classified according to the CNN. (**B**) Confusion matrix determining quantitative accuracy of the CNN for the testing set. Cell component classes include blood vessels, ducts, fat, tumor cells, wavy collagen, straight collagen, fibrotic tissue, and white space (blank space). 300 images were analyzed per class. Overall model accuracy of 93.0%. (**C**) Qualitative analysis of CNN model accuracy showing original histology images side-by-side with the CNN classified image. The first set of images highlights the model’s ability to identify blood vessels in both fat and wavy collagen (Fig. 2C,i). The second set of images recognizes the distinction of ducts, both excretory and terminal, in wavy collagen (Fig. 2C,ii). The third set of images shows the detection of cancer cells, straight collagen, and fibrotic tissue (Fig. 2C,iii). Scale bars in black are 100 μm. Color legend for each classified feature is included in the figure.

### Straight collagen strongly correlates with global stiffness

Histograms of fully classified whole-tissue slides provided cell and ECM composition for all tissue samples (Fig. 3a). Stiffness measurements of tumor-adjacent and tumor tissues revealed that both global stiffness and composition were heterogeneous within each patient (Fig. 3a). Mechanically soft tissue included the highest percentages of fat and wavy collagen (Fig. 3a). The tissues with the highest Young’s moduli contained greater percentages of blood vessels, tumor cells, straight collagen, and fibrotic tissue (Fig. 3a).

**Fig. 3.**
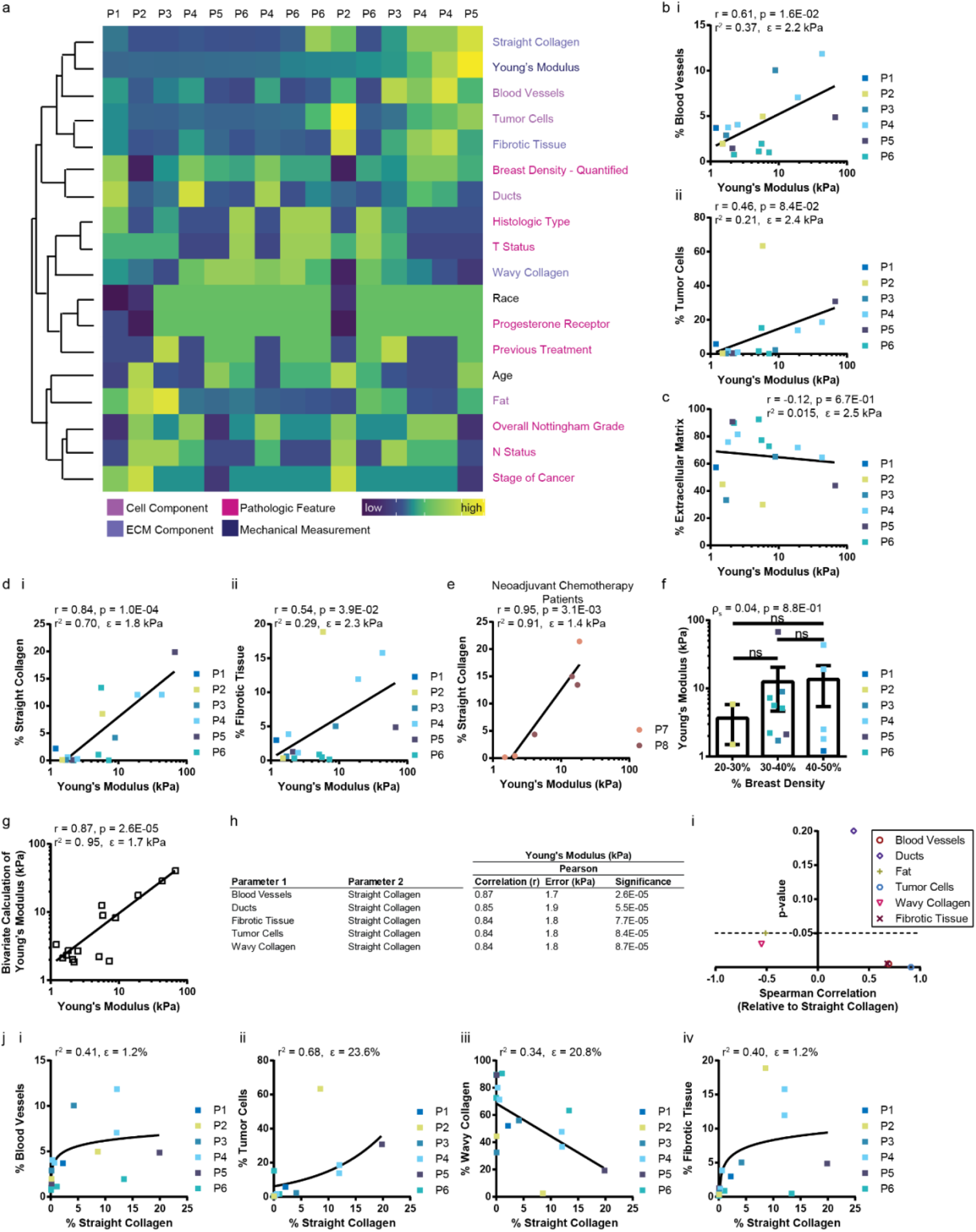
Young’s modulus (global stiffness) characterization and composition analysis of breast tissue. (**A**) Heatmap including (columns) 15 tissue samples from 6 patients (P#) clustered using Euclidean distance with complete linkage by (rows) related features. Each parameter is normalized using a z score. The values within each feature are color coded by low to high. The heatmap key on the left denotes the following color-coded parameters of each feature: cell component, extracellular matrix (ECM) component, pathologic feature, or mechanical measurement. (**B**) Univariate analysis comparing Young’s modulus (global stiffness; kPa) to the percent composition of cell component class: (**i**) blood vessels, (**ii**) tumor cells; (**C**) extracellular matrix combined; (**D**) (**i**) straight collagen and (**ii**) fibrotic tissue; (**E**) straight collagen from patients who received neoadjuvant chemotherapy; and (**F)** percent breast density. (**G**) Highest correlated pair of tissue composition classes with Young’s Modulus. The Pearson Correlation (r), p-value, r^2^ value, and error is listed at the top of plots **B-E** and **G**. One-way ANOVA was used to perform statistics in **F**. (**H**) Table of top five correlated tissue composition pairs from bivariate analysis using normal distribution and identity link using MATLAB’s glmfit and glmval functions. Rank ordered by correlation. The error is the fit-error. (**I**) Plot of Spearman Correlation (ρ_s_) versus the p-value for all cellular and extracellular classes versus straight collagen. Values below the dashed line where p=0.05 are significant. (**J**) Plots showing the monotonic relationship between straight collagen and (**i**) blood vessels, (**ii**) tumor cells, (**iii**) wavy collagen, and (**iv**) fibrotic tissue. Plots in **J** show the r^2^ value and root mean squared error (RMSE) at the top of the plot. Plots with square data points represent luminal A patients who have not received chemotherapy. Plots with circles represent patients who received neoadjuvant chemotherapy. Each data point is color coded by patient. The lines denote the best fit trend line.

Further analysis of the data suggested that the Young’s modulus, the global stiffness measurement of each tissue, has a logarithmic relationship with each tissue component.*(62, 63)* Plots of the log stiffness value versus the percent composition of each class yielded a linear line of best fit and associated Pearson correlation. Blood vessels had a significant but only moderately strong positive correlation with global stiffness (r=0.61, *p*=0.016), suggesting that this relationship was important but did not fully describe the system (Fig. 3b,i). Highlighted by the fact that tissue with the greatest tumor cell composition belonged to a tissue with a stiffness value of 5.8 kPa, while the lowest composition belonged to a stiffness of 7.2 kPa (r=0.46, *p*=0.084) (Fig. 3b,ii), tumor stiffness did not always increase with the percentage of tumor cells. Combining all matrix (non-cellular) classes into one category revealed that there was no clear correlation (r=−0.12, *p*=0.67) between the total extracellular matrix content and global breast tissue stiffness (Fig. 3c). This finding may be a result of the high percentage of wavy collagen, an ECM class that did not significantly correlate with stiffness in each tissue sample. While the percentage of fibrotic tissue showed a moderately strong correlation (r=0.54, *p*=0.039) with the Young’s modulus of the tissue (Fig. 3d,ii), there was a strong positive correlation (r=0.84, *p*=0.0001) between the percentage of straight collagen and the Young’s modulus (Fig. 3d,i). Parsing the extracellular matrix classes demonstrated that the necessity of evaluating ECM components separately from the bulk.

In the clinic, neoadjuvant chemotherapy is known to be a confounding factor in the resulting breast tissue composition as it contributes to the generation of fibrotic tissue.*(64, 65)* In two patients who received neoadjuvant chemotherapy, there was a significantly strong positive correlation (r=0.95, *p*=0.0031) between straight collagen and Young’s modulus (Fig. 3e). This result suggests that the relationship between straight collagen and global stiffness is independent of whether a patient has received neoadjuvant chemotherapy.

The eight patients in the luminal A, non-neoadjuvant chemotherapy cohort had mammographically heterogeneously dense breasts (Table 1). When quantified, this category spanned a range of 20-50% dense breast tissue (Fig. 3f, Table 1). Binning of the percent density into three categories showed that there was no significant relationship between breast density and global tissue stiffness in our study (Fig. 3f). The Spearman correlation between the two parameters was effectively zero (Fig. 3f).

A general linearized model was used to perform bivariate analysis of tissue composition classes in the patients without neoadjuvant chemotherapy (see methods). The stiffness measurements were converted into log scale values prior to running the analysis. The correlation between Young’s Modulus and any two tissue classes only slightly increases in strength (r=0.87, *p*=0.000026) (Fig. 3g). The effect of straight collagen dominates the top five strongest bivariate correlations (Fig. 3h). The percentage of blood vessels in combination with straight collagen yielded the highest correlation (Fig. 3g-h). This result is supported by the above univariate analysis (Fig. 3b,i and 3d,i).

### Straight collagen content correlates with other cellular and extracellular classes

Given the importance of straight collagen composition in determining breast tissue stiffness, we investigated the relationship of straight collagen composition to other cellular and extracellular classes (Fig. 3i-j). Tissue stiffness is often discussed and compared based on orders of magnitude changes, and frequently visualized on a logarithmic scale.*(62, 63)* Unlike Young’s modulus, the quantitative relationship between various cellular and extracellular classes has not been extensively studied. Thus, we cannot assume that the percentage of straight collagen has a linear, proportional response with the other tissue components, and have chosen to report the Spearman correlation instead of the Pearson correlation.

There is a significant, moderately strong Spearman correlation (ρ_s_) (ρ_s_=0.69, *p*=0.0045) between the percentage of blood vessels and straight collagen (Fig. 3i). The positive correlation means that a higher percentage of blood vessels moderately parallels a higher percentage of straight collagen. The best fit line to describe the relationship was logarithmic (Fig. 3j,i). Increased vascular density has been linked to poor tumor differentiation and an increase in cancer cell proliferation,*(66)* which suggests that there may be a trade-off between vascularization and an effort by cancer cells to align collagen.

The percentage of tumor cells has a strongly positive correlation with straight collagen (ρ_s_=0.91, *p*=0.0000024) (Fig. 3i). This relationship suggests a near perfect monotonic relationship between these parameters, and agrees with our understanding of tumor biology that tumor cells are responsible for restructuring the extracellular matrix to create aligned fibers.*(42, 67, 68)* The line of best fit for the data based on the R-squared value is an exponential curve, however the root mean squared error (RMSE) is high using this fit (Fig. 3j,ii). This finding is distinct from the earlier observation that the percentage of tumor cells does not strongly or significantly correlate with tissue stiffness (ρ_s_=0.55, *p*=0.035; r=0.46, *p*=0.084) (Fig. 3b,ii). The correlations between each combination of the three parameters suggest complex relationships between tumor development through changes in tissue composition and mechanical properties like tissue stiffness.

With respect to the other ECM classes, the percentage of straight collagen increased as wavy collagen decreased (ρ_s_=−0.55, *p*=0.034) and fibrotic tissue increased (ρ_s_=0.68, *p*=0.0054) (Fig. 3i). The best fit line for wavy collagen was linear but had a high RMSE (Fig. 3j,iii). The degree of collagen curvature, i.e. straight versus curly, was previously related to its location from the tumor,*(61, 69, 70)* and found to be independent of the grade of malignancy.*(61)* For fibrotic tissue, the best fit line was logarithmic (Fig. 3j,iv).

### Local stiffness is best described by straight collagen content

Local measurements reveal the large variations in stiffness values of a fresh patient tissue sample (Fig. 4a). Manual registration of the microindentation values on to the CNN-classified histology image allowed for the direct comparison between local elastic moduli and local tissue composition (Fig. 4b). Manual registration was necessary since the microindentation images and map are performed on whole tissues lacking resolution required to identify tissue components, and therefore the section must be aligned to the whole tissue image based on whole tissue shape and knowledge of microindentation sampling. Compositions were determined for the region directly under the microindenter, i.e. a 500 μm-diameter circle (Fig. 4b,inset). Two tissue samples from one patient in the luminal A non-neoadjuvant cohort, not previously used in the global stiffness analysis, were chosen for the local stiffness analysis since the processed samples could be directly matched to the unprocessed images obtained from microindentation mapping.

**Fig. 4.**
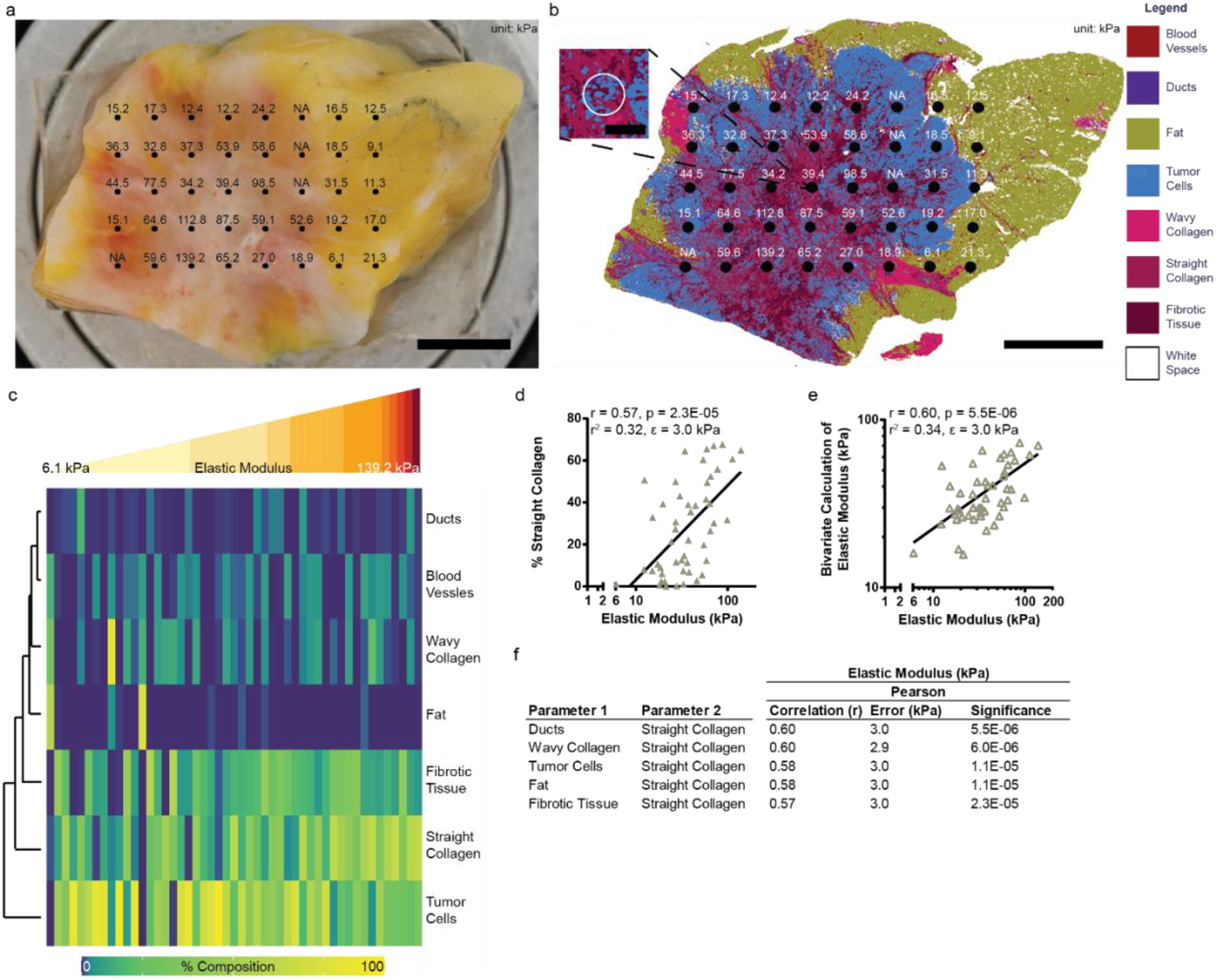
Microindentation mapping, characterization, and composition analysis of breast tissue. Fresh patient tissue with elastic modulus (local stiffness; kPa) map overlay. Scale bar in black is 5000 μm. (**B**) Corresponding Convolutional Neural Network (CNN) classified image of the patient tissue in (**A**) with the microindentation stiffness (kPa) map overlay. Scale bar in black is 5000 μm. **B inset**, Inset shows the composition of a representative microindentation point. Scale bar in black is 500 μm. Bad measurements are listed as NA and do not contribute to the analysis. (**C**) Heatmap clustered using Euclidean distance with complete linkage by (row) each cell or extracellular matrix class detailing the percent composition (0 to 100%). Each column is a different microindentation point organized from the lowest to the highest stiffness (kPa) value (49 measurements, 2 tissues, 1 patient). (**D)** Univariate analysis comparing elastic modulus (local stiffness; kPa) to the percent composition of straight collagen. (**E**) Bivariate analysis showcasing the tissue composition pair with the highest correlation to local stiffness. The line denotes the best fit line. The Pearson Correlation (r), p-value, r^2^ value, and fit-error is listed at the top of the plot. (**F**) Table highlighting the top five tissue composition pairs correlated with the elastic modulus. Rank ordered by correlation. The error is the fit-error.

Visualizing the increasing local stiffness demonstrates that the indentations with the greatest stiffness values had the highest percentages of straight collagen (Fig. 4c). The greatest percentages of tumor cells and fat coincided with some of the lower and middle stiffness values (Fig. 4c). As with the global Young’s modulus, we considered the logarithm of the local elastic modulus versus the tissue classes. The log of the elastic modulus had a significant, moderately strong linear relationship with straight collagen (r=0.57, *p*=0.000023) (Fig. 4d). This was the only cellular or extracellular relationship to the elastic modulus that was significant. Bivariate analysis only slightly increased the correlation of the tissue composition classes to the elastic modulus (r=0.60, *p*=0.0000055) (Fig. 4e). Straight collagen again dominated the top five correlations (Fig. 4f). The strongest bivariate pair is ducts combined with straight collagen (Fig. 4e-f).

### Breast density does not strongly correlate with tissue classes

The concept of breast density is often conflated with breast tissue stiffness. We showed using global stiffness measurements that quantified breast density does not have a clear correlation with Young’s modulus of the tissue (Fig. 3f). We sought an answer to the question of which cellular or extracellular classes relate to quantified breast density. The relationship between component and percentage breast density was determined using two methods. The first was through a Spearman correlation (ρ_s_), highlighting a monotonic relationship between ranked values (Fig. 5a). The second was by binning the percent breast density into three intervals and comparing the composition (Fig. 5b-h).

**Fig. 5.**
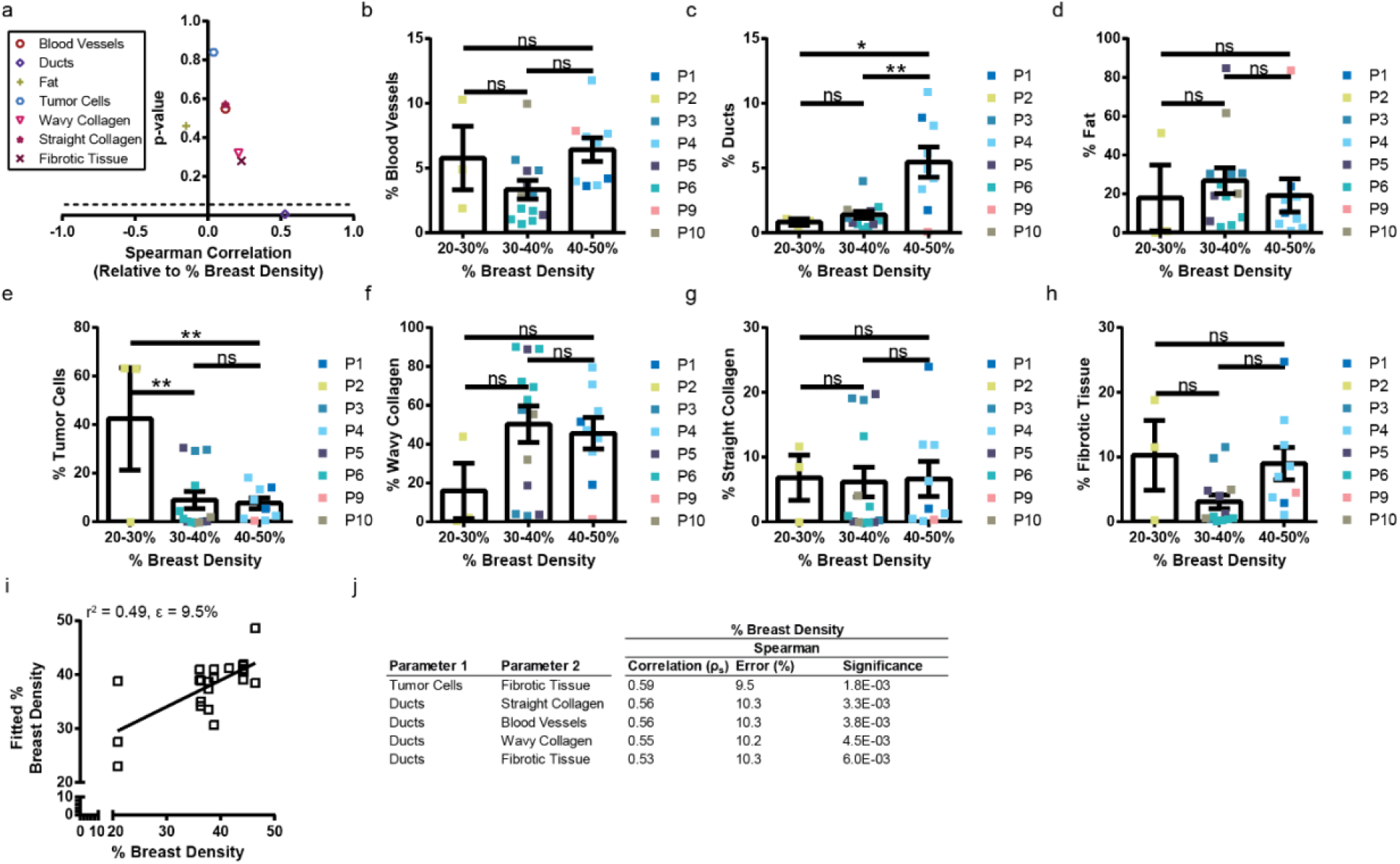
Breast density does not correlate with tissue composition. (**A**) Plot of Spearman Correlation (ρ_s_) versus the p-value for all cellular and extracellular classes versus percent breast density. Values below the dashed line where p=0.05 are significant. Percent breast density versus cell classes (**B**) blood vessels, (**C**) ducts, (**D**) fat, (**E**) tumor cells; and extracellular classes (**F**) wavy collagen, (**G**) straight collagen, (**H**) fibrotic tissue. When binned, the quantified breast density is related via bar chart using a one-way ANOVA. (**I**) Bivariate analysis showing the highest pair of features that correlate with the percent breast density. The r^2^ value and fit-error are at the top of the plot. The line denotes the best fit line. (**J**) Table highlighting the top five tissue composition pairs correlated with the percent of breast density. Rank ordered by correlation. The error is the fit-error.

The percentage of blood vessels and fat alone did not correlate with percent density (Fig. 5a) and was not significantly different from 20-50% dense breast tissue (Fig. 5b). The percentage of ducts had a significant, moderately positive correlation with percent of dense breast tissue (Fig. 5a). Furthermore, 40-50% dense breast tissue has significantly greater percentage of ducts than 20-30% or 30-40% dense breast tissue (Fig. 5c). While this initial finding is in line with the current understanding that dense breast tissue highlights an increase in glandular tissue,*(2)* future work spanning a larger patient population (i.e. wider range of breast densities, more patients with lower mammographic breast densities) is necessary to validate this claim. Even though we did not find a correlation between tumor cells alone and breast density (ρ_s_ = 0.04, *p* = 0.84) (Fig. 5a), there were more tumor cells in tissues with a breast density of 20-30% than 30-40% and 40-50% (Fig. 5e). There was no significant correlation between any of the extracellular matrix classes and the percent breast density (Fig. 5a), nor was there a significant relationship between breast densities within each of the components (Fig 5f-h). Assuming a normal distribution, tumor cells and fibrotic tissue combined had a significant and moderately positive correlation with breast density (ρ_s_=0.59, *p*=0.0018) (Fig. 5i-j). After this first combination, the percentage of ducts dominated the bivariate relation (Fig. 5j).

## Discussion

Through tissue component identification using a deep learning model, we were able to connect mechanical measurements to patient tissue composition. We identified the highest univariate correlate of both global and local stiffness to be straight collagen (Fig. 3d,i and Fig. 4d). Our findings improve upon and depart from previous work with these key discoveries: straight collagen is a biomechanical marker in human tissue; straight collagen has strong monotonic relationships with other cellular and extracellular classes; Young’s modulus is dependent on tissue composition; the fibrillar phenotype is identifiable using H&E without SHG or additional staining; and that straight collagen does not directly relate to breast density. Furthermore, we use whole tissue slides in our analysis affording us both the ability to analyze a greater fraction of each tumor than tissue microarrays (TMAs), and to utilize the same slides already procured in the clinic for diagnostics and treatment. Previously, through the use of TMAs, straightened and aligned collagen was linked to poor disease-specific and disease-free survival independent of the cancer type, stage of cancer, hormone status, and node status.*(71)* Aligned collagen perpendicular to the tumor acts as a mechanism for local invasion by cancer cells.*(68, 72)* This class was proposed as a predictor for breast cancer survival, i.e. that increased aligned collagen suggests poor prognosis.*(71, 73)* In mice, the elastic modulus of mammary glands was shown to increase in tumors due to collagen crosslinking, which then forms more fibrillar and aligned collagen.*(42)* In *ex vivo* measurements of human breast tissues, the mean Young’s modulus was shown to vary based on the tissue and histologic tumor type.*(74)* In an earlier attempt to relate breast tissue stiffness and breast density, the stiffness was approximated from a theoretical calculation of breast volume and area in a mammogram, not through actual mechanical measurements of the tissue.*(75)*

Treatment of all stroma as a single class would have led to the incorrect conclusion that the extracellular matrix does not contribute to mechanical stiffness in patient tissue (Fig. 3c). The predominance of the wavy collagen phenotype, which on average accounts for 56.6% of the classes identified in each tissue section, causes this misleading result when the ECM is bundled into a single category. Our results following separation of wavy and straight fibrillar collagen and fibrotic tissue highlights the importance of separating ECM classes into pathologically relevant subtypes. Future analysis, through immunohistochemical and immunofluorescent staining, could incorporate the identification of immune and stromal cells.

The relationship between tumor cells, straight collagen, and Young’s Modulus is worth briefly discussing as these classes defining stiffer tumor sections are in line with the current understanding of tumor biology.*(29, 76–79)* Tumor cells have a weak correlation with stiffness, despite their strong correlation with straight collagen. We think that there is a difference between the maximum stiffness achievable by a cell component versus that of an extracellular class. Tumor cells can align collagen but are not stiff themselves; therefore, the aligned collagen has a greater contribution to tissue stiffness.

All patients used in the study of breast density had a luminal A subtype and were designated as having categorical heterogeneously dense breasts. Within this specific category, the quantified breast density ranged between 20-50%. In contrast to previously reported literature, we did not find that mammographic density correlated with aligned collagen.*(80, 81)* The different findings could be a result of earlier works identifying low and high mammographic density from a patient cohort with values predominantly below 20% density,*(80)* or with patients across multiple breast density categories.*(81)*

Our result urges caution when discussing breast density versus breast stiffness. Additionally, this outcome supports the clinically accepted separation between findings from palpations and cancer occurrence or prognosis.*(82–84)* Of note, the tissue samples are from regions in or near the excised tumor region and may not fully represent the non-excised regions of the breast, whereas the breast density determination is based on the whole breast. We are unable to specifically trace back the excised tissue sample to an exact area of high or low mammographic density in the mammogram image. Future studies would need to know the exact location of the excised tissue to relate the tissue composition findings to regions of breast tissue density in mammograms, and utilize a larger patient cohort with a range of categorical and quantitative breast density. Furthermore, while we did relate breast density to both the Young’s modulus for global tissue stiffness and the elastic modulus for local tissue stiffness, there are other types of stiffness measurements that could have distinct relationships with breast density.

## Materials and Methods

### Patient tissues

Patients with abnormal screening or diagnostic breast imaging findings require pathologic examination (either core needle aspiration or less frequently fine needle aspiration) to definitively characterize the abnormal radiographic lesion. If positive for breast cancer, the pathologist will determine the histologic subtype, assign a Nottingham histologic grade, and perform additional breast biomarker studies (Fig. 1a). The combination of physical examination and imaging modalities help to assign the clinical staging regarding the size of the tumor (T), abnormal axillary lymph node (N) and the presence of metastatic disease (M). If the patient undergoes surgical resection, lumpectomy or mastectomy, the pathological staging will be reported by the size of the mass (T) and any lymph node involvement (N). During the pathologic evaluation, the histologic type and Nottingham score are confirmed, and the overall pathology cancer stage is assigned as defined by the American Joint Committee of Cancer Staging Manual, 8^th^ edition*(57)* (Primary Tumor [T] Status and Regional Lymph Nodes [N] Status) (Fig. 1a).

All patient tissue samples were obtained with written consent from the patient and approved by the Johns Hopkins Medicine Institutional Review Board (IRB). Tumor-adjacent and tumor tissue samples received from the patients were kept in 4°C DPBS immediately after mastectomy or lumpectomy. Tumor samples were then transferred for mechanical tests within 4 h of resection. The tumor tissue was then sectioned to expose the regions of interest for micromechanical mapping and bulk compression tests.

Fifteen tissues from six luminal A patients that did not receive neoadjuvant chemotherapy were chosen for the global stiffness analysis. Six tissues from two patients, one with luminal A subtype and one with TNBC subtype, that received neoadjuvant chemotherapy were used in a separate analysis of the relationship between global stiffness and tissue composition to avoid any confounding tissue composition distributions associated with neoadjuvant chemotherapy previously reported in the literature.*(25, 64, 85)* Two tissues from one patient with a luminal A subtype and no neoadjuvant chemotherapy were used for complementary local stiffness analysis. Only luminal A patients who did not receive neoadjuvant chemotherapy were used to analyze quantified breast density. Tissue samples from all patients were used to train the neural network.

### Microindentation of tissues

The tumor section was mounted on a customized stage and DPBS was applied to keep the tissue hydrated throughout the measurement. Dynamic indentation by a nanoindenter (Nanomechanics Inc.) was used to characterize the tumor elastic modulus.*(86)* Sneddon’s stiffness equation*(87)* was applied to relate dynamic stiffness of the contact to the elastic storage modulus of the samples.*(88, 89)* 500 μm flat cylindrical probe was used in the indentation experiments. Briefly, procedure of indentation is comprised of 3 steps: 1) approaching and finding tissue surface at the indenter’s resonant frequency to enhance contact sensitivity and accuracy, 2) pre-compression of 50 μm to ensure good contact, 3) dynamic measurement at 100Hz oscillation frequency with amplitude of 250 nm. The indentation procedure mentioned above was done consecutively on multiple regions of a single tissue surface in a grid pattern to obtain elastic moduli map of the tumor. Because obtaining a perfectly flat tissue surface was difficult due to tissue heterogeneity, individual indentation processes were observed using a microscope camera to determine inappropriate contact of the probe to the tissue for inaccurate measurement which were excluded from data. Typically, the number of indentation points per tissue mapping was 20-40 with the resolution of 2.5 ± 0.5 mm spacing between points depending on the size of tumor sample. The duration of stiffness mapping was 30 min on average, not exceeding 40 min. A single measurement was obtained for each indentation.

### Compression test of tissues

Tissue samples were sectioned to obtain flat and parallel surfaces on all sides. Once the sample was sectioned, it was immediately staged on tensile/compression tester (MTS Criterion) for measurement.*(90)* Top compression plate was lowered until in full contact with tissue sample at minimal load. Once in contact, the samples could relax and stabilize for 1 min before actual compression test. Tissue samples were compressed at 0.25 mm/sec deformation rate until 20% strain. Young’s modulus calculation was done on the best-fitted slope of the initial linear region (~5-10%) of the obtained stress-strain curve. A single measurement was obtained for each tissue.

### Patient tissue processing

After obtaining mechanical measurements, each tissue was fixed in formalin for 24 h. The tissue was transferred to PBS prior to embedding in paraffin, sectioning (4 μm), and staining with hematoxylin and eosin (H&E). To minimize the batch effects of H&E image staining and scanning conditions, all tissues were stained in and scanned by the same laboratory.

### Quantifying breast density from mammograms

Pectoral muscle was removed from mammogram images prior to receipt. Images were then cropped to remove any identifiers and keep only the breast image. The image was then converted to type 8-bit. Thresholding was performed using MinError(l) in ImageJ and a histogram was taken to determine the total breast pixel size. Reverting to the original 8-bit image, thresholding using Moments and taking a histogram determined the number of dense breast tissue pixels. A breast density percentage was obtained by dividing the number of white pixels from the Moments thresholding by the number of white pixels using MinError(l) thresholding and multiplying by 100.

### Second-harmonic generation

Mounted tissue slides were imaged using a LD LCI Plan-Apochromat 25x/0.8 Imm objective mounted on a Zeiss LSM 710 NLO upright microscope. Excitation was provided by a Chameleon Vision II mode-locked Ti:Sapphire laser tuned to 880 nm, and the SHG signal was captured by an epi-mounted non-descanned detector with a 420-480nm bandpass filter.

### Manual annotations

Manual annotations of tissue slides were performed using Aperio ImageScope [v12.3.3.5048]. Briefly, cellular and extracellular components were identified manually in H&E-stained tissue slides by outlining the feature using the built-in annotation function. Within each tissue slide, we annotated 30 or more instances of a feature type to create the tissue and non-tissue-based classes. The annotations were verified by a trained pathologist.

### Convolutional neural network architecture

We used H&E stained slides of breast tumor-adjacent and tumor tissues to train the CNN.*(60)* The slides were scanned at 20x, with a spatial resolution of 0.5μm/pixel, and down-sampled using the openslide library*(91)* to a pixel size of 1μm/pixel. Example regions of different tissue classes were manually annotated (30+ annotations per tissue class) in each individual slide. In this study, we annotated seven tissue classes including blood vessels, ducts, fat, tumor cells, wavy collagen, straight collagen, and fibrotic tissue; and one non-tissue class which we term white space. The CNN was trained and validated in MATLAB 2019b with 3600 randomly selected non-repeating image tiles per annotation class from all patient slides. Of these 3600 images per class, 3000 were used for training, and 300 were used for validation and testing. Dropout layers and a window size of 103 pixels × 103 pixels × 3 channels were used to facilitate the classification of both cellular and extracellular classes in the model. The training images were augmented via positive or negative 90° rotations to increase the training size and prevent overfitting.*(92–95)* Adam (adaptive moment estimation) optimization was used with an initial learning rate of 0.013 to train the model. Training finished when validation accuracy did not improve for five epochs. The network architecture of the CNN model contains four convolutional layers each followed by a batch normalization and rectified linear unit (ReLu) layers. The second convolutional layer is followed by a dropout layer of 0.1. Then there are six convolutional layers in parallel, each with a batch and ReLu layer. An additional layer and ReLu layer are added before five more convolutional/batch/ReLu layers. There is a max pooling layer, convolutional layer, dropout layer of 0.1, batch and ReLu layers. Next, a convolutional/batch/ReLu/max pooling set before a fully connected layer with batch normalization and ReLu layers. The architecture ends with a fully connected layer, batch normalization layer, and softmax output layer.

### Computation of tissue composition

Classified images were imported into ImageJ. Histogram analysis of the whole tissue section provided tissue composition values for global stiffness (15 tissue samples, 6 patients). For local stiffness composition, the fresh patient tissue image contains the original microindentation map overlay. The CNN classified image was scaled and manually registered to match the original fresh patient tissue image. Histogram analysis inside of 500 μm (62.5 px) diameter circles on the CNN classified image provided the local stiffness composition (3 tissue samples, 2 patients).

### Bivariate and univariate analysis

MATLAB’s built-in function ‘corr’ was used to perform univariate analysis resulting in either a Pearson or Spearman correlation and statistical significance. MATLAB’s built-in functions ‘glmfit’ and ‘glmval’ were used to perform bivariate analysis resulting in a correlation coefficient, fit error, and statistical significance for each pair. The global and local stiffness measurements are converted to log base 10 values before analysis. The distribution used was ‘normal,’ and the link was ‘identity.’ The general form of the equation is:

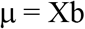

where μ is the response with a normal distribution, X is a matrix of predictors, and b is a vector of coefficient estimates.

The number of patient tissue samples and patients for each parameter are as follows: global stiffness – 15 tissue samples, 6 patients; local stiffness – 2 tissue samples, 1 patient; breast density quantification – 20 tissue samples, 8 patients.

### Heatmaps of tissue composition, mechanical measurements, and pathologic features

Heatmaps of global and local stiffness data were created in RStudio using R version 3.6.3 and function superheat. Clustering was performed using Euclidean distance with a complete linkage method.

### Statistical analysis

Statistical analysis for univariate and bivariate analysis plots and tables was performed using MATLAB’s “corr” function. The line of best fit was plotted using Prism 6 (GraphPad Software, Inc.). For the breast density bar chart analysis, ordinary one-way ANOVAs using Turkey’s multiple comparison test with a single pooled variance were performed in Prism 6 (GraphPad Software, Inc.). All bar chart graphs are reported as mean ± SEM. * p < 0.05, **p < 0.01, *** p< 0.001, and *** p< 0.0001.

## Supplementary Materials

Fig. S1. Comparison of H&E tissue features with CNN classified image.

Fig. S2. Non-significant relationships between tissue composition and Young’s modulus (global stiffness).

## Acknowledgments

We would like to thank Dr. Tatiana Prowell, Dr. Eti Cukierman, and Dr. Mary Daly for their insight into breast density, histology, clinical outcomes, and mechanobiology. We thank the Johns Hopkins Oncology Tissue Service for embedding, sectioning, and staining the tissues. We also thank Ryan O’Connor and Muna Igboko for their help with manual annotations of the patient tissues.

## Author contributions

A.S. formulated the project, performed manual annotations, created the deep learning dataset, trained the network, analyzed the data, managed the project, and wrote the manuscript. J.K. and L.G. performed mechanical testing on the patient tissue. A.K. created the deep learning software and assisted in its implementation. M.W. reviewed the H&E annotations, and pathologic data. P.W. and J.M.P. aided in the analysis. M.H. provided the primary patient tissue. D.W. formulated the project, managed the project, and wrote the manuscript. A.S., J.K., A.K., M.H., M.W., P.W. J.M.P., L.G., and D.W. read and edited the manuscript.

## Data and materials availability

The code for the deep learning package, bivariate and univariate analysis will be made available on GitHub upon publication. The datasets that support the findings of this study are available from the corresponding author upon request.

## References and Notes

1. D. Wirtz, K. Konstantopoulos, P. C. Searson, The physics of cancer: The role of physical interactions and mechanical forces in metastasis, Nat. Rev. Cancer 11, 512–522 (2011).

2. Centers for Disease Control and Prevention, What Does It Mean to Have Dense Breasts? | CDC (2020) (available at https://www.cdc.gov/cancer/breast/basic_info/dense-breasts.htm).

3. T. M. Kolb, J. Lichy, J. H. Newhouse, Comparison of the performance of screening mammography, physical examination, and breast US and evaluation of factors that influence them: An analysis of 27,825 patient evaluations, Radiology 225, 165–175 (2002).

4. S. S. Nazari, P. Mukherjee, An overview of mammographic density and its association with breast cancerBreast Cancer 25, 259–267 (2018).

5. S. M. Conroy, I. Pagano, L. N. Kolonel, G. Maskarinec, Mammographic density and hormone receptor expression in breast cancer: The Multiethnic Cohort Study, Cancer Epidemiol. 35, 448–452 (2011).

6. N. F. Boyd, J. M. Rommens, K. Vogt, V. Lee, J. L. Hopper, M. J. Yaffe, A. D. Paterson, Mammographic breast density as an intermediate phenotype for breast cancerLancet Oncol. 6, 798–808 (2005).

7. D. B. Kopans, Basic physics and doubts about relationship between mammographically determined tissue density and breast cancer riskRadiology 246, 348–353 (2008).

8. N. F. Boyd, H. Guo, L. J. Martin, L. Sun, J. Stone, E. Fishell, R. A. Jong, G. Hislop, A. Chiarelli, S. Minkin, M. J. Yaffe, Mammographic Density and the Risk and Detection of Breast Cancer, N. Engl. J. Med. 356, 227–236 (2007).

9. T. L. Nguyen, Y. K. Aung, S. Li, N. H. Trinh, C. F. Evans, L. Baglietto, K. Krishnan, G. S. Dite, J. Stone, D. R. English, Y.-M. Song, J. Sung, M. A. Jenkins, M. C. Southey, G. G. Giles, J. L. Hopper, Predicting interval and screen-detected breast cancers from mammographic density defined by different brightness thresholds, Breast Cancer Res. 20, 152 (2018).

10. N. Boyd, H. Berman, J. Zhu, L. J. Martin, M. J. Yaffe, S. Chavez, G. Stanisz, G. Hislop, A. M. Chiarelli, S. Minkin, A. D. Paterson, The origins of breast cancer associated with mammographic density: A testable biological hypothesis, Breast Cancer Res. 20(2018), doi:10.1186/s13058-018-0941-y.

11. A. Burton, G. Maskarinec, B. Perez-Gomez, C. Vachon, H. Miao, M. Lajous, R. López-Ridaura, M. Rice, A. Pereira, M. L. Garmendia, R. M. Tamimi, K. Bertrand, A. Kwong, G. Ursin, E. Lee, S. A. Qureshi, H. Ma, S. Vinnicombe, S. Moss, S. Allen, R. Ndumia, S. Vinayak, S. H. Teo, S. Mariapun, F. Fadzli, B. Peplonska, A. Bukowska, C. Nagata, J. Stone, J. Hopper, G. Giles, V. Ozmen, M. E. Aribal, J. Schüz, C. H. Van Gils, J. O. P. Wanders, R. Sirous, M. Sirous, J. Hipwell, J. Kim, J. W. Lee, C. Dickens, M. Hartman, K. S. Chia, C. Scott, A. M. Chiarelli, L. Linton, M. Pollan, A. A. Flugelman, D. Salem, R. Kamal, N. Boyd, I. dos-Santos-Silva, V. McCormack, Mammographic density and ageing: A collaborative pooled analysis of cross-sectional data from 22 countries worldwide, PLoS Med. 14(2017), doi:10.1371/journal.pmed.1002335.

12. C. Byrne, G. Ursin, C. F. Martin, J. D. Peck, E. B. Cole, D. Zeng, E. Kim, M. D. Yaffe, N. F. Boyd, G. Heiss, A. Mctiernan, R. T. Chlebowski, D. S. Lane, J. E. Manson, J. Wactawski-Wende, E. D. Pisano, Mammographic Density Change With Estrogen and Progestin Therapy and Breast Cancer Risk, J Natl Cancer Inst 109(2017), doi:10.1093/jnci/djx001.

13. A. Burton, G. Byrnes, J. Stone, R. M. Tamimi, J. Heine, C. Vachon, V. Ozmen, A. Pereira, M. L. Garmendia, C. Scott, J. H. Hipwell, C. Dickens, J. Schüz, M. E. Aribal, K. Bertrand, A. Kwong, G. G. Giles, J. Hopper, B. Pérez Gómez, M. Pollán, S.-H. Teo, S. Mariapun, N. A. M. Taib, M. Lajous, R. Lopez-Riduara, M. Rice, I. Romieu, A. A. Flugelman, G. Ursin, S. Qureshi, H. Ma, E. Lee, R. Sirous, M. Sirous, J. W. Lee, J. Kim, D. Salem, R. Kamal, M. Hartman, H. Miao, K.-S. Chia, C. Nagata, S. Vinayak, R. Ndumia, C. H. van Gils, J. O. P. Wanders, B. Peplonska, A. Bukowska, S. Allen, S. Vinnicombe, S. Moss, A. M. Chiarelli, L. Linton, G. Maskarinec, M. J. Yaffe, N. F. Boyd, I. dos-Santos-Silva, V. A. McCormack, Mammographic density assessed on paired raw and processed digital images and on paired screen-film and digital images across three mammography systems, Breast Cancer Res. 18, 130 (2016).

14. S. Elsamany, A. Alzahrani, S. A. Elkhalik, O. Elemam, E. Rawah, M. U. Farooq, M. H. Almatrafi, F. K. Olayan, Prognostic value of mammographic breast density in patients with metastatic breast cancer, Med. Oncol. 31, 1–7 (2014).

15. T. Cil, E. Fishell, W. Hanna, P. Sun, E. Rawlinson, S. A. Narod, D. R. McCready, Mammographic density and the risk of breast cancer recurrence after breast-conserving surgery, Cancer 115, 5780–5787 (2009).

16. S. Y. H. Chiu, S. Duffy, A. M. F. Yen, L. Tabár, R. A. Smith, H. H. Chen, Effect of baseline breast density on breast cancer incidence, stage, mortality, and screening parameters: 25-Year follow-up of a Swedish mammographic screening, Cancer Epidemiol. Biomarkers Prev. 19, 1219–1228 (2010).

17. C. H. Van Gils, J. D. M. Otten, A. L. M. Verbeek, J. H. C. L. Hendriks, R. Holland, Effect of mammographic breast density on breast cancer screening performance: A study in Nijmegen, the Netherlands, J. Epidemiol. Community Health 52, 267–271 (1998).

18. C. Colpaert, P. Vermeulen, E. Van Marck, L. Dirix, The presence of a fibrotic focus is an independent predictor of early metastasis in lymph node-negative breast cancer patients., Am. J. Surg. Pathol. 25, 1557–8 (2001).

19. T. Li, L. Sun, N. Miller, T. Nicklee, J. Woo, L. Hulse-Smith, M.-S. Tsao, R. Khokha, L. Martin, N. Boyd, The Association of Measured Breast Tissue Characteristics with Mammographic Density and Other Risk Factors for Breast Cancer, Cancer Epidemiol Biomarkers Prev 14, 343–349 (2005).

20. S. M. Thompson, J. Wang, V. S. Chandan, K. J. Glaser, L. R. Roberts, R. L. Ehman, S. K. Venkatesh, MR elastography of hepatocellular carcinoma: Correlation of tumor stiffness with histopathology features—Preliminary findings, Magn. Reson. Imaging 37, 41–45 (2017).

21. A. Evans, P. Whelehan, K. Thomson, D. McLean, K. Brauer, C. Purdie, L. Jordan, L. Baker, A. Thompson, Quantitative shear wave ultrasound elastography: Initial experience in solid breast masses, Breast Cancer Res. 12, R104 (2010).

22. M. Denis, A. Gregory, M. Bayat, R. T. Fazzio, D. H. Whaley, K. Ghosh, S. Shah, M. Fatemi, A. Alizad, Correlating tumor stiffness with immunohistochemical subtypes of breast cancers: Prognostic value of comb-push ultrasound shear elastography for differentiating luminal subtypes, PLoS One 11(2016), doi:10.1371/journal.pone.0165003.

23. J. M. Chang, I. A. Park, S. H. Lee, W. H. Kim, M. S. Bae, H. R. Koo, A. Yi, S. J. Kim, N. Cho, W. K. Moon, Stiffness of tumours measured by shear-wave elastography correlated with subtypes of breast cancer, Eur. Radiol. 23, 2450–2458 (2013).

24. S. Imtiaz, Breast elastography : A New paradigm in diagnostic breast imaging, Appl. Radiol. 47, 14–19 (2018).

25. M. Hayashi, Y. Yamamoto, M. Ibusuki, S. Fujiwara, S. Yamamoto, S. Tomita, M. Nakano, K. Murakami, K. I. Iyama, H. Iwase, Evaluation of tumor stiffness by elastography is predictive for pathologic complete response to neoadjuvant chemotherapy in patients with breast cancer, Ann. Surg. Oncol. 19, 3042–3049 (2012).

26. A. Thomas, S. Kümmel, F. Fritzsche, M. Warm, B. Ebert, B. Hamm, T. Fischer, Real-Time Sonoelastography Performed in Addition to B-Mode Ultrasound and Mammography: Improved Differentiation of Breast Lesions?, Acad. Radiol. 13, 1496–1504 (2006).

27. H. Zhi, B. Ou, B. M. Luo, X. Feng, Y. L. Wen, H. Y. Yang, Comparison of ultrasound elastography, mammography, and sonography in the diagnosis of solid breast lesions, J. Ultrasound Med. 26, 807–815 (2007).

28. M. L. Bennett, C. J. Welman, L. M. Celliers, How reassuring is a normal breast ultrasound in assessment of a screen-detected mammographic abnormality? A review of interval cancers after assessment that included ultrasound evaluation, Clin. Radiol. 66, 928–939 (2011).

29. J. G. Goetz, S. Minguet, I. Navarro-Lérida, J. J. Lazcano, R. Samaniego, E. Calvo, M. Tello, T. Osteso-Ibáñez, T. Pellinen, A. Echarri, A. Cerezo, A. J. P. Klein-Szanto, R. Garcia, P. J. Keely, P. Sánchez-Mateos, E. Cukierman, M. A. Del Pozo, Biomechanical remodeling of the microenvironment by stromal caveolin-1 favors tumor invasion and metastasis, Cell 146, 148–163 (2011).

30. K. M. Wisdom, K. Adebowale, J. Chang, J. Y. Lee, S. Nam, R. Desai, N. S. Rossen, M. Rafat, R. B. West, L. Hodgson, O. Chaudhuri, Matrix mechanical plasticity regulates cancer cell migration through confining microenvironments, Nat. Commun. 9, 1–13 (2018).

31. Y. A. Miroshnikova, G. I. Rozenberg, L. Cassereau, M. Pickup, J. K. Mouw, G. Ou, K. L. Templeman, E.-I. Hannachi, K. J. Gooch, A. L. Sarang-Sieminski, A. J. García, V. M. Weaver, α5β1-Integrin promotes tension-dependent mammary epithelial cell invasion by engaging the fibronectin synergy site., Mol. Biol. Cell 28, 2958–2977 (2017).

32. M. Nebuloni, L. Albarello, A. Andolfo, C. Magagnotti, L. Genovese, I. Locatelli, G. Tonon, E. Longhi, P. Zerbi, R. Allevi, A. Podestà, L. Puricelli, P. Milani, A. Soldarini, A. Salonia, M. Alfano, Insight on Colorectal Carcinoma Infiltration by Studying Perilesional Extracellular Matrix, Sci. Rep. 6, 1–13 (2016).

33. B. R. Seo, P. Bhardwaj, S. Choi, J. Gonzalez, R. C. A. Eguiluz, K. Wang, S. Mohanan, P. G. Morris, B. Du, X. K. Zhou, L. T. Vahdat, A. Verma, O. Elemento, C. A. Hudis, R. M. Williams, D. Gourdon, A. J. Dannenberg, C. Fischbach, Obesity-dependent changes in interstitial ECM mechanics promote breast tumorigenesis, Sci. Transl. Med. 7, 301ra130–301ra130 (2015).

34. A. Parekh, N. S. Ruppender, K. M. Branch, M. K. Sewell-Loftin, J. Lin, P. D. Boyer, O. E. Candiello, W. D. Merryman, S. A. Guelcher, A. M. Weaver, Sensing and modulation of invadopodia across a wide range of rigidities, Biophys. J. 100, 573–582 (2011).

35. R. G. Wells, The role of matrix stiffness in regulating cell behaviorHepatology 47, 1394–1400 (2008).

36. Y. H. Bae, K. L. Mui, B. Y. Hsu, S. L. Liu, A. Cretu, Z. Razinia, T. Xu, E. Puré, R. K. Assoian, A FAK-Cas-Rac-lamellipodin signaling module transduces extracellular matrix stiffness into mechanosensitive cell cycling, Sci. Signal. 7, ra57 (2014).

37. J. Schrader, T. T. Gordon-Walker, R. L. Aucott, M. van Deemter, A. Quaas, S. Walsh, D. Benten, S. J. Forbes, R. G. Wells, J. P. Iredale, Matrix stiffness modulates proliferation, chemotherapeutic response, and dormancy in hepatocellular carcinoma cells, Hepatology 53, 1192–1205 (2011).

38. A. Pathak, S. Kumar, Independent regulation of tumor cell migration by matrix stiffness and confinement, Proc. Natl. Acad. Sci. U. S. A. 109, 10334–10339 (2012).

39. J. K. Mouw, Y. Yui, L. Damiano, R. O. Bainer, J. N. Lakins, I. Acerbi, G. Ou, A. C. Wijekoon, K. R. Levental, P. M. Gilbert, E. S. Hwang, Y. Y. Chen, V. M. Weaver, Tissue mechanics modulate microRNA-dependent PTEN expression to regulate malignant progression, Nat. Med. 20, 360–367 (2014).

40. H. Yu, J. K. Mouw, V. M. Weaver, Forcing form and function: biomechanical regulation of tumor evolution, (2010), doi:10.1016/j.tcb.2010.08.015.

41. M. J. Paszek, N. Zahir, K. R. Johnson, J. N. Lakins, G. I. Rozenberg, A. Gefen, C. A. Reinhart-King, S. S. Margulies, M. Dembo, D. Boettiger, D. A. Hammer, V. M. Weaver, Tensional homeostasis and the malignant phenotype, Cancer Cell 8, 241–254 (2005).

42. K. R. Levental, H. Yu, L. Kass, J. N. Lakins, M. Egeblad, J. T. Erler, S. F. T. Fong, K. Csiszar, A. Giaccia, W. Weninger, M. Yamauchi, D. L. Gasser, V. M. Weaver, Matrix crosslinking forces tumor progression by enhancing integrin signaling., Cell 139, 891–906 (2009).

43. M. Gilkes, G. L. Semenza, D. Wirtz, Hypoxia and the extracellular matrix: drivers of tumour metastasis, Nat. Rev. Cancer 14, 430–439 (2014).

44. O. M. T. Pearce, R. M. Delaine-Smith, E. Maniati, S. Nichols, J. Wang, S. Böhm, V. Rajeeve, D. Ullah, P. Chakravarty, R. R. Jones, A. Montfort, T. Dowe, J. Gribben, J. L. Jones, H. M. Kocher, J. S. Serody, B. G. Vincent, J. Connelly, J. D. Brenton, C. Chelala, P. R. Cutillas, M. Lockley, C. Bessant, M. M. Knight, F. R. Balkwill, Deconstruction of a metastatic tumor microenvironment reveals a common matrix response in human cancers, Cancer Discov. 8, 304–319 (2018).

45. L. Neary-Zajiczek, C. Essmann, N. Clancy, A. Haider, E. Miranda, M. Shaw, A. Gander, B. Davidson, D. Fernandez-Reyes, V. Pawar, D. Stoyanov, in Lecture Notes in Computer Science, D. Shen, et al., Eds. (Springer, 2019), vol. 11764 LNCS, pp. 760–768.

46. M. Plodinec, M. Loparic, C. A. Monnier, E. C. Obermann, R. Zanetti-Dallenbach, P. Oertle, J. T. Hyotyla, U. Aebi, M. Bentires-Alj, R. Y. H. Lim, C. A. Schoenenberger, The nanomechanical signature of breast cancer, Nat. Nanotechnol. 7, 757–765 (2012).

47. R. Thavarajah, V. K. Mudimbaimannar, J. Elizabeth, U. K. Rao, K. Ranganathan, Chemical and physical basics of routine formaldehyde fixationJ.Oral Maxillofac. Pathol. 16, 400–405 (2012).

48. R. G. Wells, Tissue mechanics and fibrosisBiochim. Biophys. Acta - Mol. Basis Dis. 1832, 884–890 (2013).

49. A. Dolor, F. C. Szoka, Digesting a Path Forward: The Utility of Collagenase Tumor Treatment for Improved Drug Delivery, Mol. Pharm. 15, 2069–2083 (2018).

50. S. Amar, G. B. Fields, Potential clinical implications of recent matrix metalloproteinase inhibitor design strategiesExpertRev. Proteomics 12, 445–447 (2015).

51. G. Baumgartner, C. Gomar-Höss, L. Sakr, E. Ulsperger, C. Wogritsch, in Cancer Letters, (Elsevier, 1998), vol. 131, pp. 85–99.

52. B. Benson, Z. A. Wainberg, J. R. Hecht, D. Vyushkov, H. Dong, J. Bendell, F. Kudrik, A Phase II Randomized, Double-Blind, Placebo-Controlled Study of Simtuzumab or Placebo in Combination with Gemcitabine for the First-Line Treatment of Pancreatic Adenocarcinoma, Oncologist 22, 241 (2017).

53. S. R. Hingorani, A. J. Bullock, T. E. Seery, L. Zheng, D. Sigal, P. S. Ritch, F. S. Braiteh, M. Zalupski, N. Bahary, W. P. Harris, J. Pu, C. Aldrich, S. Khelifa, X. W. Wu, J. Baranda, P. Jiang, E. Hendifar, Randomized phase II study of PEGPH20 plus nab-paclitaxel/gemcitabine (PAG) vs AG in patients (Pts) with untreated, metastatic pancreatic ductal adenocarcinoma (mPDA)., J. Clin. Oncol. 35, 4008–4008 (2017).

54. H. Ma, J. Luo, M. F. Press, Y. Wang, L. Bernstein, G. Ursin, Is there a difference in the association between percent mammographic density and subtypes of breast cancer? luminal a and triple-negative breast cancer, Cancer Epidemiol. Biomarkers Prev. 18, 479–485 (2009).

55. E. Mema, F. Schnabel, J. Chun, E. Kaplowitz, A. Price, J. Goodgal, L. Moy, The relationship of breast density in mammography and magnetic resonance imaging in women with triple negative breast cancer, Eur. J. Radiol. 124(2020), doi:10.1016/j.ejrad.2020.108813.

56. S. AJ, M. M, S. S, N. R, A. I, O. J, W. R, M. Z, G. GL, Mammographic Breast Density and Breast Cancer Molecular Subtypes: The Kenyan-African Aspect, Biomed Res. Int. 2018(2018), doi:10.1155/2018/6026315.

57. M. Amin, S. Edge, F. Greene, D. Byrd, R. Brookland, et al. Washington, MK, Gershenwald JE, Compton CC, Hess KR, AJCC Cancer Staging Manual (Springer International Publishing: American Joint Commission on Cancer, ed. 8th, 2017).

58. J. N. Kather, A. T. Pearson, N. Halama, D. Jäger, J. Krause, S. H. Loosen, A. Marx, P. Boor, F. Tacke, U. P. Neumann, H. I. Grabsch, T. Yoshikawa, H. Brenner, J. Chang-Claude, M. Hoffmeister, C. Trautwein, T. Luedde, Deep learning can predict microsatellite instability directly from histology in gastrointestinal cancer, Nat. Med. 25, 1054–1056 (2019).

59. D. F. Steiner, R. MacDonald, Y. Liu, P. Truszkowski, J. D. Hipp, C. Gammage, F. Thng, L. Peng, M. C. Stumpe, Impact of Deep Learning Assistance on the Histopathologic Review of Lymph Nodes for Metastatic Breast Cancer, Am. J. Surg. Pathol. 42, 1636–1646 (2018).

60. A. Kiemen, A. M. Braxton, M. P. Grahn, K. S. Han, J. M. Babu, R. Reichel, F. Amoa, S. Hong, T. C. Cornish, E. D. Thompson, L. D. Wood, R. H. Hruban, P.-H. Wu, D. Wirtz, In situ characterization of the 3D microanatomy of the pancreas and pancreatic cancer at single cell resolution, bioRxiv (2020), doi:https://doi.org/10.1101/2020.12.08.416909.

61. A. Brabrand, I. I. Kariuki, M. J. Engstrøm, O. A. Haugen, L. A. Dyrnes, B. O. Åsvold, M. B. Lilledahl, A. M. Bofin, Alterations in collagen fibre patterns in breast cancer. A premise for tumour invasiveness?, APMIS 123, 1–8 (2015).

62. J. W. Shin, D. J. Mooney, Extracellular matrix stiffness causes systematic variations in proliferation and chemosensitivity in myeloid leukemias, Proc. Natl. Acad. Sci. U. S. A. 113, 12126–12131 (2016).

63. H. C. Wells, K. H. Sizeland, S. J. R. Kelly, N. Kirby, A. Hawley, S. Mudie, R. G. Haverkamp, Collagen Fibril Intermolecular Spacing Changes with 2-Propanol: A Mechanism for Tissue Stiffness, (2017), doi:10.1021/acsbiomaterials.7b00418.

64. S. Wu, Y. Huang, Q. Tang, Z. Li, H. Horng, J. Li, Z. Wu, Y. Chen, H. Li, Quantitative evaluation of redox ratio and collagen characteristics during breast cancer chemotherapy using two-photon intrinsic imaging, Biomed. Opt. Express 9, 1375 (2018).

65. H. Honkoop, H. M. Pinedo, J. S. De Jong, H. M. W. Verheul, S. C. Linn, K. Hoekman, J. Wagstaff, P. J. Van Diest, Effects of Chemotherapy on Pathologic and Biologic Characteristics of Locally Advanced Breast Cancer, Am. J. Clin. Pathol. 107(1997) (available at https://academic.oup.com/ajcp/article-abstract/107/2/211/1756800).

66. P. Lipponen, H. Ji, S. Aaltomaa, K. Syrjänen, Tumour vascularity and basement membrane structure in breast cancer as related to tumour histology and prognosis, J. Cancer Res. Clin. Oncol. 120, 645–650 (1994).

67. M. Egeblad, M. G. Rasch, V. M. Weaver, Dynamic interplay between the collagen scaffold and tumor evolutionCurr. Opin. Cell Biol. 22, 697–706 (2010).

68. P. P. Provenzano, K. W. Eliceiri, J. M. Campbell, D. R. Inman, J. G. White, P. J. Keely, Collagen reorganization at the tumor-stromal interface facilitates local invasion, BMC Med. 4, 38 (2006).

69. S. Pearson, C. J. Hall, C. R. Reid, G. Falzon, Small-angle X-ray scattering and second-harmonic generation imaging studies of collagen in invasive carcinoma Cell tracking with x-rays and nano particles View project X-ray Fluorescence Computed Tomography View project, Am. Inst. Phys. (2006) (available at https://www.researchgate.net/publication/233809700).

70. G. Falzon, S. Pearson, R. Murison, Analysis of collagen fibre shape changes in breast cancer, Phys. Med. Biol 53, 6641–6652 (2008).

71. M. W. Conklin, J. C. Eickhoff, K. M. Riching, C. A. Pehlke, K. W. Eliceiri, P. P. Provenzano, A. Friedl, P. J. Keely, Aligned Collagen Is a Prognostic Signature for Survival in Human Breast Carcinoma, Am. J. Pathol. 178, 1221–1232 (2011).

72. I. Acerbi, L. Cassereau, I. Dean, Q. Shi, A. Au, C. Park, Y. Y. Chen, J. Liphardt, E. S. Hwang, V. M. Weaver, Human breast cancer invasion and aggression correlates with ECM stiffening and immune cell infiltration, Integr. Biol. (United Kingdom) 7, 1120–1134 (2015).

73. J. Bredfeldt, Y. Liu, M. Conklin, P. Keely, T. Mackie, K. Eliceiri, Automated quantification of aligned collagen for human breast carcinoma prognosis, J. Pathol. Inform. 5, 28 (2014).

74. A. Samani, J. Zubovits, D. Plewes, Elastic moduli of normal and pathological human breast tissues: an inversion-technique-based investigation of 169 samples, Phys. Med. Biol. 52, 1565–1576 (2007).

75. N. F. Boyd, Q. Li, O. Melnichouk, E. Huszti, L. J. Martin, A. Gunasekara, G. Mawdsley, M. J. Yaffe, S. Minkin, J. R. Rees, Ed. Evidence That Breast Tissue Stiffness Is Associated with Risk of Breast Cancer, PLoS One 9, e100937 (2014).

76. D. Fukumura, D. G. Duda, L. L. Munn, R. K. Jain, Tumor Microvasculature and Microenvironment: Novel Insights Through Intravital Imaging in Pre-Clinical Models, Microcirculation 17, 206–225 (2010).

77. T. D. Tlsty, L. M. Coussens, Tumor Stroma and Regulation of Cancer Development, Annu. Rev. Pathol. Mech. Dis. 1, 119–150 (2006).

78. P. P. Provenzano, K. W. Eliceiri, P. J. Keely, Multiphoton microscopy and fluorescence lifetime imaging microscopy (FLIM) to monitor metastasis and the tumor microenvironmentClin. Exp. Metastasis 26, 357–370 (2009).

79. M. W. Conklin, J. C. Eickhoff, K. M. Riching, C. A. Pehlke, K. W. Eliceiri, P. P. Provenzano, A. Friedl, P. J. Keely, Aligned collagen is a prognostic signature for survival in human breast carcinoma, Am. J. Pathol. (2011), doi:10.1016/j.ajpath.2010.11.076.

80. J. C. McConnell, O. V. O’Connell, K. Brennan, L. Weiping, M. Howe, L. Joseph, D. Knight, R. O’Cualain, Y. Lim, A. Leek, R. Waddington, J. Rogan, S. M. Astley, A. Gandhi, C. C. Kirwan, M. J. Sherratt, C. H. Streuli, Increased peri-ductal collagen micro-organization may contribute to raised mammographic density, Breast Cancer Res. 18, 1–17 (2016).

81. C. W. Huo, G. Chew, P. Hill, D. Huang, W. Ingman, L. Hodson, K. A. Brown, A. Magenau, H. Allam, E. McGhee, P. Timpson, M. A. Henderson, E. W. Thompson, K. Britt, High mammographic density is associated with an increase in stromal collagen and immune cells within the mammary epithelium, Breast Cancer Res. 17, 1–20 (2015)

82. .A. K. Hackshaw, E. A. Paul, Breast self-examination and death from breast cancer: A meta-analysis, Br. J. Cancer 88, 1047–1053 (2003).

83. J. P. Kösters, P. C. Gøtzsche, Regular self-examination or clinical examination for early detection of breast cancer., Cochrane database Syst. Rev., CD003373 (2003).

84. L. WW, C. CP, C. CF, M. CC, C. KW, L. MH, T. MH, Factors affecting the palpability of breast lesion by self-examination., Singapore Med. J. 49, 228–232 (2008).

85. H. AH, P. HM, D. J. JS, V. HM, L. SC, H. K, W. J, van D. PJ, Effects of Chemotherapy on Pathologic and Biologic Characteristics of Locally Advanced Breast Cancer, Am. J. Clin. Pathol. 107(1997), doi:10.1093/AJCP/107.2.211.

86. R. Akhtar, E. R. Draper, D. J. Adams, J. Hay, Oscillatory nanoindentation of highly compliant hydrogels: A critical comparative analysis with rheometry, J. Mater. Res. 33, 873–883 (2018).

87. N. Sneddon, The relation between load and penetration in the axisymmetric boussinesq problem for a punch of arbitrary profile, Int. J. Eng. Sci. 3, 47–57 (1965).

88. E. G. Herbert, W. C. Oliver, A. Lumsdaine, G. M. Pharr, Measuring the constitutive behavior of viscoelastic solids in the time and frequency domain using flat punch nanoindentation, J. Mater. Res. 24, 626–637 (2009).

89. E. G. Herbert, W. C. Oliver, G. M. Pharr, Nanoindentation and the dynamic characterization of viscoelastic solids, J. Phys. D. Appl. Phys. 41(2008), doi:10.1088/0022-3727/41/7/074021.

90. G. N. Grifno, A. M. Farrell, R. M. Linville, D. Arevalo, J. H. Kim, L. Gu, P. C. Searson, Tissue-engineered blood-brain barrier models via directed differentiation of human induced pluripotent stem cells, Sci. Rep. 9, 1–13 (2019).

91. D. Forsberg, fordanic/openslide-matlab (2020) (available at https://www.github.com/fordanic/openslide-matlab).

92. A. Gertych, Z. Swiderska-Chadaj, Z. Ma, N. Ing, T. Markiewicz, S. Cierniak, H. Salemi, S. Guzman, A. E. Walts, B. S. Knudsen, Convolutional neural networks can accurately distinguish four histologic growth patterns of lung adenocarcinoma in digital slides, Sci. Rep. 9, 1–12 (2019).

93. A. Serag, A. Ion-Margineanu, H. Qureshi, R. McMillan, M. J. Saint Martin, J. Diamond, P. O’Reilly, P. Hamilton, Translational AI and Deep Learning in Diagnostic PathologyFront. Med. 6(2019), doi:10.3389/fmed.2019.00185.

94. H. Reza Tizhoosh, L. Pantanowitz, Artificial intelligence and digital pathology: Challenges and opportunities, J. Pathol. Inform. 9(2018), doi:10.4103/jpi.jpi_53_18.

95. D. Komura, S. Ishikawa, Machine Learning Methods for Histopathological Image AnalysisComput. Struct. Biotechnol. J. 16, 34–42 (2018).

