## Supplementary Figures for "Deep Learning Identification of Stiffness Markers in Breast Cancer"

**Affiliations:**
[a] *Department of Chemical and Biomolecular Engineering, Johns Hopkins Physical Sciences-Oncology Center, and Institute for NanoBioTechnology, Johns Hopkins University, 3400 N Charles St, Baltimore, Maryland 21218, USA.*

[b] *Department of Materials Science and Engineering, and Institute for NanoBioTechnology, Johns Hopkins University, 3400 N Charles St, Baltimore, Maryland 21218, USA.*

[c] *Johns Hopkins Breast Center, Johns Hopkins Bayview Medical Center, 4940 Eastern Ave, Baltimore, Maryland 21224, USA.*

[d] *Department of Pathology, Johns Hopkins School of Medicine, 401 N Broadway, Baltimore, Maryland 21231, USA.*

[e] *Department of Biomedical Engineering, Johns Hopkins University, 3400 N Charles St, Baltimore, Maryland 21218, USA.*

[f] *Department of Oncology, Johns Hopkins School of Medicine, 1800 Orleans St, Baltimore, Maryland 21205, USA.*

**Supplementary Materials:**

**Fig. S1**. Comparison of H&E tissue features with CNN classified image.

**Fig. S2**. Non-significant relationships between tissue composition and Young's modulus (global stiffness).

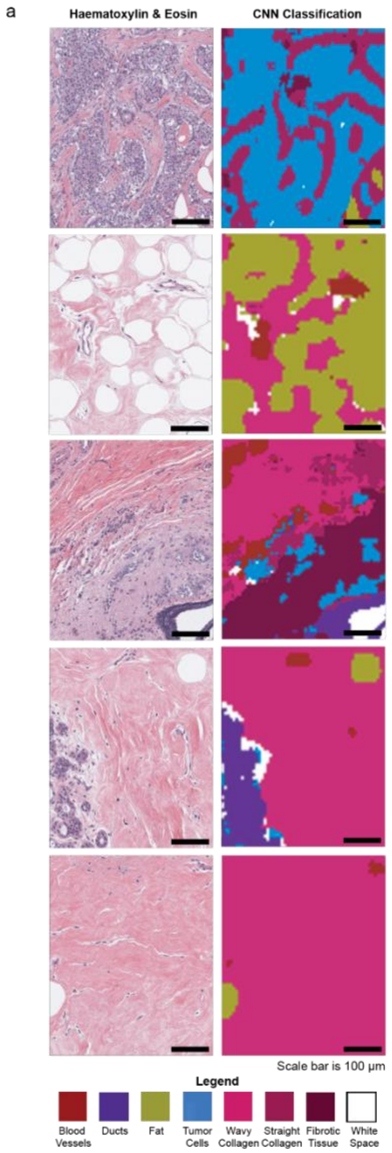


**Fig. S1**. Comparison of H&E tissue features with CNN classified image. (**A**) Qualitative analysis of CNN model accuracy showing original histology images side-by-side with the CNN classified image. Scale bars in black are 100 µm. Color legend for each classified feature is included in the figure.

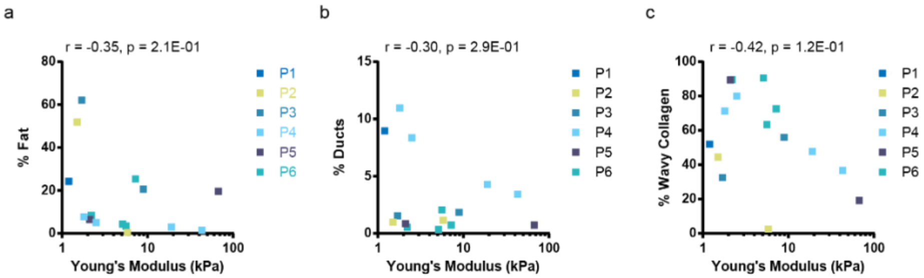


**Fig. S2.** Non-significant relationships between tissue composition and Young's modulus (global stiffness). Univariate analysis comparing Young's modulus (global stiffness; kPa) to the percent composition of cell component class (**A**) fat, (**B**) ducts; and extracellular matrix class (**C**) wavy collagen. The Pearson Correlation (r) and p-value are listed at the top of plots **A-C**.
